# Manipulation of the Nuclear Envelope-Associated Protein SLAP During Mammalian Brain Development Affects Cortical Lamination and Exploratory Behavior

**DOI:** 10.1101/2024.01.31.578288

**Authors:** Ivan Mestres, Azra Atabay, Joan-Carles Escolano, Solveig Arndt, Klara Schmidtke, Maximilian Einsiedel, Melina Patsonis, Lizbeth Airais Bolanos Castro, Maximina Yun, Nadine Bernhardt, Anna Taubenberger, Federico Calegari

## Abstract

Here we report the first characterization of the effects resulting from the manipulation of Soluble-Lamin Associated Protein (SLAP) expression during mammalian brain development. We found that SLAP localizes to the nuclear envelope and when overexpressed causes changes in nuclear morphology and lengthening of mitosis. SLAP overexpression in apical progenitors of the developing mouse brain altered asymmetric cell division, neurogenic commitment and neuronal migration ultimately resulting in unbalance in the proportion of upper, relative to deeper, neuronal layers. Several of these effects were also recapitulated upon Cas9-mediated knock-down. Ultimately, SLAP overexpression during development resulted in a reduction in subcortical projections of young mice and, notably, reduced their exploratory behavior. Our study shows the potential relevance of the previously uncharacterized nuclear envelope protein SLAP in neurodevelopmental disorders.

## INTRODUCTION

Mammalian brain development relies on the correct balance between proliferation and differentiation of neural stem cells, as well as specification and migration of newborn neurons to achieve proper cortical lamination. Defects in these processes often cause developmental defects and cognitive impairment (Parenti et al., 2020).

In the embryonic cortex, neural stem cells reside within the ventricular zone (VZ) lining the ventricle. Their division can be symmetric to amplify the stem cell pool, asymmetric to generate neurogenic progenitors forming the subventricular zone (SVZ) or, to a lesser degree, neurons directly (Agirman et al., 2017). In turn, post-mitotic neurons migrate from the germinal zones across the intermediate zone (IZ) to reach their final destination within the cortical plate (CP) (Lui et al., 2011; Taverna et al., 2014). Of the various organelles and sub-cellular compartments that are known to play fundamental roles in coordinating neural stem cell fate and corticogenesis, the nuclear envelope (NE) has received the least attention.

The NE consists of two lipid bilayers within which nuclear pore complexes are embedded that mediate the exchange between the nucleus and the cytoplasm. Additionally, intermediate filaments called lamins are anchored to the inner nuclear membrane forming a mesh that provides physical support and serves as a scaffold for NE-associated proteins (Hetzer, 2010). The NE modulates several fundamental processes. For example, it separates the genome from the cytoplasm, couples the nucleus to the cytoskeleton for nuclear and cell migration, and participates in chromatin remodeling to control gene expression (Balaji et al., 2022; Hetzer, 2010; Kalukula et al., 2022). Furthermore, the NE disassembles during mitosis and reassembles in daughter cells, leading to defective chromosome segregation and DNA damage when not properly coordinated (Dauer and Worman, 2009). While these aspects are common to most cell types, few studies has focused on how components of the NE participate in the differentiation of somatic stem cells in general and neural stem fate commitment in particular.

Among the NE-associated proteins, the nuclear pore component Nup133 recruits the motor protein dynein to modulate neural stem cell nuclear migration and cell cycle progression during corticogenesis (Hu et al., 2013) as well as gene expression during adult neurogenesis (Toda et al., 2017). In addition, mutant mice for several NE-associated proteins, including Sun1/2, Syne2 and Tor1A, exhibited several neurodevelopmental defects such as progenitor depletion, failed neuronal migration, and defects in cortical lamination; which in some cases translated into behavioral impairments (McCarthy et al., 2012; Tanabe et al., 2016; Tsai et al., 2020; Zhang et al., 2009). Several reports focused on lamins, perhaps the most studied components of the NE during neural stem cell fate. For example, lamin B1/B2 knockout during mammalian brain development led to altered spindle orientation and asymmetric cell division of neural progenitors together with nuclear abnormalities and cortical layering defects (Coffinier et al., 2011; Kim et al., 2011). Similar defects were also observed in the retina (Razafsky et al., 2016). Notably, a decline in lamin B1 was functionally associated with depletion of neural stem cells during ageing (Bedrosian et al., 2021; bin Imtiaz et al., 2021). Indicative of their specific role in brain function, human mutations in lamin B1 were associated with microcephaly (Cristofoli et al., 2020), mainly due to a weakened nuclear integrity that resulted in defects in interkinetic nuclear migration and spindle orientation of neural progenitors as well as impaired neuronal migration and survival (Coffinier et al., 2011; Kim et al., 2011; Vergnes et al., 2004). Conversely, duplication of lamin B1 causes leukodystrophy (Padiath et al., 2006), conceivably as a result of increased nuclear stiffness (Ferrera et al., 2014) and/or repression of oligodendrocyte genes (Heng et al., 2013). In addition, increased levels of lamin B1 found in Huntington’s disease patients were correlated with chromatin rearrangements and transcriptional dysregulation (Alcalá-Vida et al., 2021). Thus, several lines of evidence indicate that levels of NE-associated proteins must be tightly regulated in a cell type-specific manner for proper brain development.

An attempt to discover novel lamin-associated proteins resulted in the identification of 11 uncharacterized genes among which FAM169a was one of the most prominent. Subsequently named Soluble Lamin-Associated Protein (Roux et al., 2012), SLAP was found to be highly expressed in the mouse and human nervous system (Cao et al., 2019; Chen et al., 2022) and among the top 20% of genes most intolerant to functional genomic mutations suggesting a high association with neurodevelopmental disorders (Fadista et al., 2017; Karczewski et al., 2020; Shohat et al., 2017). Consistently, a frameshift mutation in the SLAP locus impaired the survival and growth of human pluripotent stem cells (Yilmaz et al., 2018), and a similar mutation was found in an autistic patient (De Rubeis et al., 2014). Moreover, a study on monozygotic twins reported SLAP among the genes whose upregulation correlated with lower intellectual scores (Yu et al., 2012). Despite these associations, the function of SLAP during brain development remains unknown.

Here we show that SLAP is critical to maintain nuclear integrity and that alterations in its expression influence neural stem cell fate as well as migration and molecular identity of newborn neurons. Combined, these effects resulted in abnormal cortical layering and a reduction in the exploratory behavior of young animals. Together, our study highlights the importance of NE integrity in general, and the role of the NE-associated protein SLAP in particular, during mammalian brian development.

## RESULTS

### SLAP overexpression disrupts nuclear morphology and delays mitosis

We began our study by a general assessment of the effects of SLAP overexpression in HEK293 cells. After validating the localization of endogenously expressed SLAP within the NE by immunocytochemistry (Fig. 1A), we investigated the effects of its overexpression. We found that increased levels of SLAP triggered abnormal nuclear morphology and numbers including micronuclei and ruptures of the NE with chromatin everting into the cytoplasm (Fig. 1B). In control, GFP-overexpressing cells only 6.0 ± 0.8% exhibited abnormal nuclei, but this number increased more than 4-fold (25.1 ± 1.7%; p < 0.01) after SLAP overexpression (Fig. 1C). Given the function of NE-associated proteins in providing physical support (Hetzer, 2010), we next used atomic force microcopy (AFM) to assess nuclear stiffness finding that SLAP overexpressing nuclei displayed significantly decreased apparent Young’s moduli, hence were less stiff compared to that of GFP transfected cells (GFP = 0.65 ± 0.02 vs SLAP = 0.44 ± 0.01 KPa, p < 0.001) (Fig. 1D).

**Figure 1.**
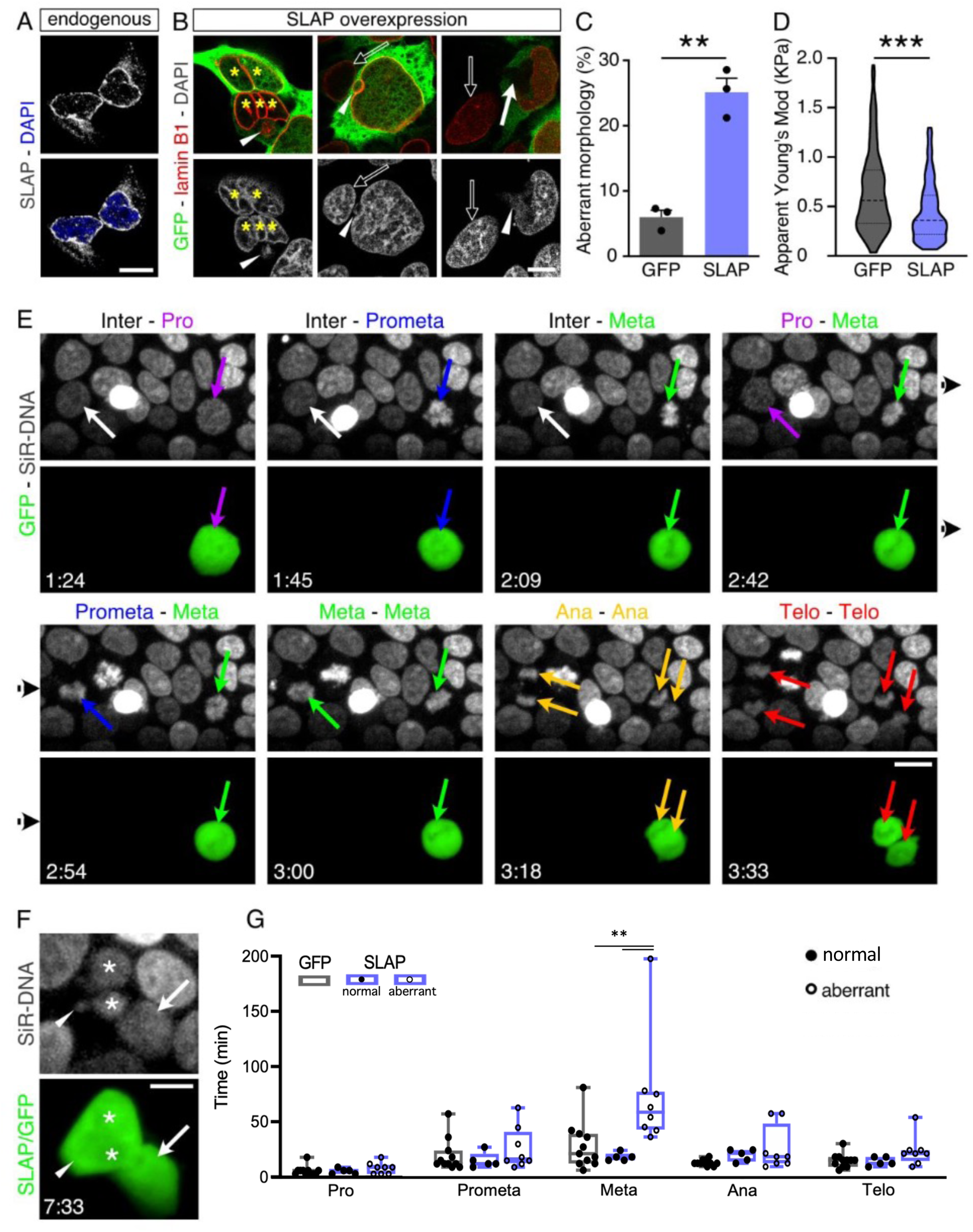
SLAP overexpression disrupts nuclear morphology and lengthens mitosis. (A and B) Immunolabeling showing endogenous SLAP (A) or lamin B1 upon SLAP overexpression together with GFP (B) in HEK239 cells. Note micronuclei (arrowheads), ruptured nucleus (arrow) and multinucleated cells (asterisks) in SLAP overexpressing cells relative to untransfected controls (empty arrows). (C and D) Quantification of nuclear aberrant morphology (C) or apparent Young’s modulus (D) in HEK293 transfected cells. (E) Snapshots of time-lapse imaging showing a SLAP transfected (green, right) and untrasfected (left) cell within the same field. Color-coded arrows indicate mitotic phases. (F) Daughters resulting from a SLAP overexpressing cell (E; green) showing nuclear fragmentation (asterisks) and a micronucleus (arrowhead) 4 h after mitosis. (G) Quantification of mitotic phases length. Filled or empty circles denote daughter cells with healthy or aberrant nuclei, respectively, as assessed > 2 h after mitosis of the mother cell. Chromatin was counterstained with DAPI (A, B) or SiR-DNA (E, F). Quantifications are depicted as mean ± SEM bar graphs (C), violin plots (D) or box plots (G). Either a two-tailed Student t test (C), a Mann-Whitney test (D), or a two-way ANOVA and Bonferroni’s post-hoc test (G) were used to assess significance (** p < 0.01, *** p < 0.001). Scale bars = 10 μm (A, B, F), 20 μm (E).

To observe how aberrant nuclear morphology arose, we transfected GFP or SLAP as above and 48 h later performed time-lapse microcopy for additional 9 h (Fig. 1E). In doing so, we focused our attention on cells undergoing mitosis and looking at their daughter cells for the appearance of nuclear abnormalities upon cytokinesis. We found not only that SLAP overexpression increased the overall duration of mitosis by about 50% (GFP = 84.7 ± 8.0 vs SLAP = 130.4 ± 12.5 min, p < 0.001; sum of the values shown in Fig. 1G, see below) but also that in only about half of the cases daughter cells started to display nuclear abnormalities 2 h after cytokinesis (Fig. 1F, G and supplementary Video 1). Hence, we next attempted to identify i) the mitotic phase(s) displaying a delay upon SLAP overexpression and ii) whether a correlation existed in mitotic delay and the generation of daughter cells with nuclear abnormalities. Strikingly, we observed that SLAP overexpression resulted in a lengthening specifically of metaphase and that such lengthening was strongly predictive of the appearance of nuclear abnormalities in daughter cells. In fact, and suggestive of a causal relationship, mitosis length of SLAP overexpressing cells resulting in daughter with normal nuclei were undistinguishable from GFP controls (Fig. 1G).

Altogether, these results show that SLAP is an important player in the regulation of mitosis, specifically metaphase, and that changes in its expression result in alterations of nuclear biomechanical properties and structure.

### SLAP overexpression favors asymmetric cell division and generation of basal progenitors

We next investigated whether SLAP overexpression triggered similar effects in the developing cortex as observed in HEK293 cells. To this end, we in utero electroporated embryonic day (E) 13 mouse embryos with either a vector expressing a cytoplasmic GFP alone or in combination with SLAP and collected the brains 2 days later.

When analyzing apical progenitors lining the ventricular border and immunolabeled with lamin B1 to identify their nuclei, we noticed that while a small fraction (2.8 ± 0.3%) of control GFP+ cells exhibited nuclear invaginations and micronuclei, this number increased 8-fold (22.4 ± 1.0%) after SLAP overexpression (Fig. 2A, B). Furthermore, electron microscopy of electroporated brains revealed that several nuclei of SLAP transfected cells, identified by GFP gold-immunolabeling, contained deep invaginations (Fig. 2C) that in some remarkable cases extended across the entire nucleus (Fig. 2D and supplementary Video 2). Also, and consistent with the longer mitosis observed in cell culture, phospho-histone 3 (PH3) immunolabeling revealed a 2-fold increase in the proportion of cells undergoing mitosis upon SLAP overexpression (GFP = 4.6 ± 0.4 vs SLAP = 8.7 ± 0.8%, p < 0.01) (Fig. 2E). Such increase in the proportion of PH3+ cells (4.1%) fits remarkably well with the observed delay in mitosis of HEK293 cells by about 40 min (Fig. 1G) when such lengthening is expressed as a proportion (4.7%) of the length of the cell cycle of apical progenitors at this developmental stage (ca. 14 hours) (Calegari et al., 2005) and suggesting that lengthening of mitosis alone is sufficient to account for the increase in mitotic figures observed after SLAP overexpression.

**Figure 2.**
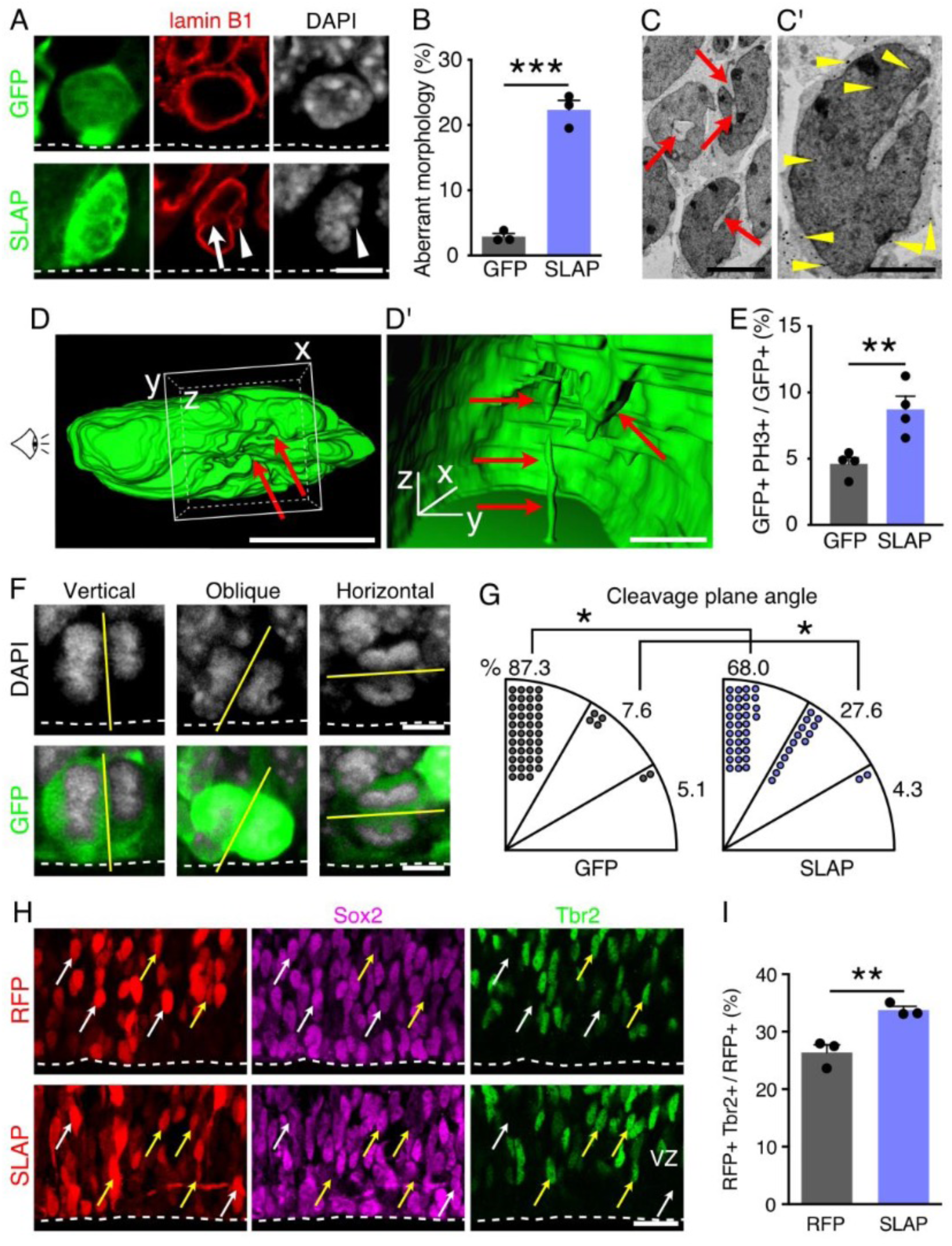
SLAP overexpression favors asymmetric division and neuronal commitment. (A and B) Images (A) of electroporated apical progenitors counterstained with lamin B1 and scored for nuclear abnormalities (B). (C and D) Electron micrographs and 3D reconstruction, with magnified insets (C’ and D’), of nuclei of GFP immunogold labeled (arrowheads) SLAP overexpressing cells showing invaginations (arrows). Drawing of eye in D indicate the point of view in D’. (E-G) Abundance (E), images (F) and proportion of cleave plane angles (G) of apical progenitors undergoing mitosis. (H and I) Images (H) of the VZ of electroporated brains counterstained with Sox2 (magenta) and Tbr2 (green) to label apical (white arrows) and basal (yellow arrows) progenitors, respectively; and quantification of the latter (I). Two-tailed Student t tests (B, E, I) or two-way ANOVA and Bonferroni’s post-hoc test (G) were used to assess significance (* p < 0.05, ** p < 0.01, *** p < 0.001). Quantifications are depicted as mean ± SEM bar graphs (B, E) or pie charts (G). Scale bars = 5 μm (A, C, D, F), 2.5 μm (C’), 1 μm (D’), 20 μm (H).

Interestingly, differences in nuclear morphology and proportion of mitotic cells upon SLAP overexpression also correlated with a change in cleavage plane orientation during anaphase with an increase in oblique, asymmetric cell divisions (30-60°: GFP = 7.6 ± 2.9 vs SLAP = 27.6 ± 4.4%, p < 0.05) (Fig. 2F, G). Considering that a lengthening of mitosis, particularly metaphase, has been reported to promote asymmetric cell division (Mora-Bermúdez et al., 2016; Pilaz et al., 2016) and that asymmetric cell division of apical progenitors is known to trigger neurogenic commitment and generation of basal progenitors (Konno et al., 2008; Kosodo et al., 2004; Noctor et al., 2004), we next performed SLAP overexpression at E13 and 2 days later assessed the proportion of Sox2+ and/or Tbr2+ cells within the VZ to identify apical and basal progenitors, respectively. To more reliably count individual cells, this time we used a nuclear-localized RFP reporter observing a minor, although significant, decrease in the proportion of Sox2+ cells (RFP = 73.6 ± 1.0% vs. SLAP = 66.5 ± 0.5%, p < 0.001) that was mirrored by an increase in the proportion of Tbr2+ basal progenitors by about one third (RFP = 26.3 ± 1.0 vs. SLA P = 33.4 ± 0.5%, p < 0.001) (Fig. 2H, I).

In essence, these experiments extended observations on HEK293 cells upon SLAP overexpression in the context of mammalian corticogenesis, including correlations between aberrant nuclear morphology, lengthening of mitosis, asymmetric cell division, neurogenic fate commitment, and generation of basal progenitors.

### SLAP overexpression alters neuronal migration, specification and cortical layering

To investigate additional effects of SLAP overexpression on neuronal cell types, we next assessed the distribution of electroporated cells across cortical layers. Two days after electroporation at E13 we noticed in SLAP overexpressing brains an increase in the proportion of cells in the VZ (RFP = 27.7 ± 1.3 vs. SLAP = 40.5 ± 2.0%, p < 0.001) and SVZ, although the latter did not reach significance (RFP = 9.7 ± 0.3 vs. SLAP = 14.8 ± 0.3%; p = 0.1), which was accompanied by a significant reduction in the CP by about 4-fold (RFP = 23.2 ± 0.8 vs. SLAP = 6.0 ± 0.6%, p < 0.001) (Fig. 3A, B). Such increase in the proportion of progenitors in the VZ/SVZ and decrease in neurons in the CP may seem in contradiction with the increase in asymmetric cell divisions and generation of neurogenic, basal progenitors described above (Fig. 2F-I). Still, these effects could be explained by additional factors including, among others, change in the survival and/or migration of newborn neurons. To evaluate these possibilities, we performed a series of immunolabelings combined with cellular birth-dating by a single administration of EdU 24 h after in utero electroporation and 24 h before sacrifice.

**Figure 3.**
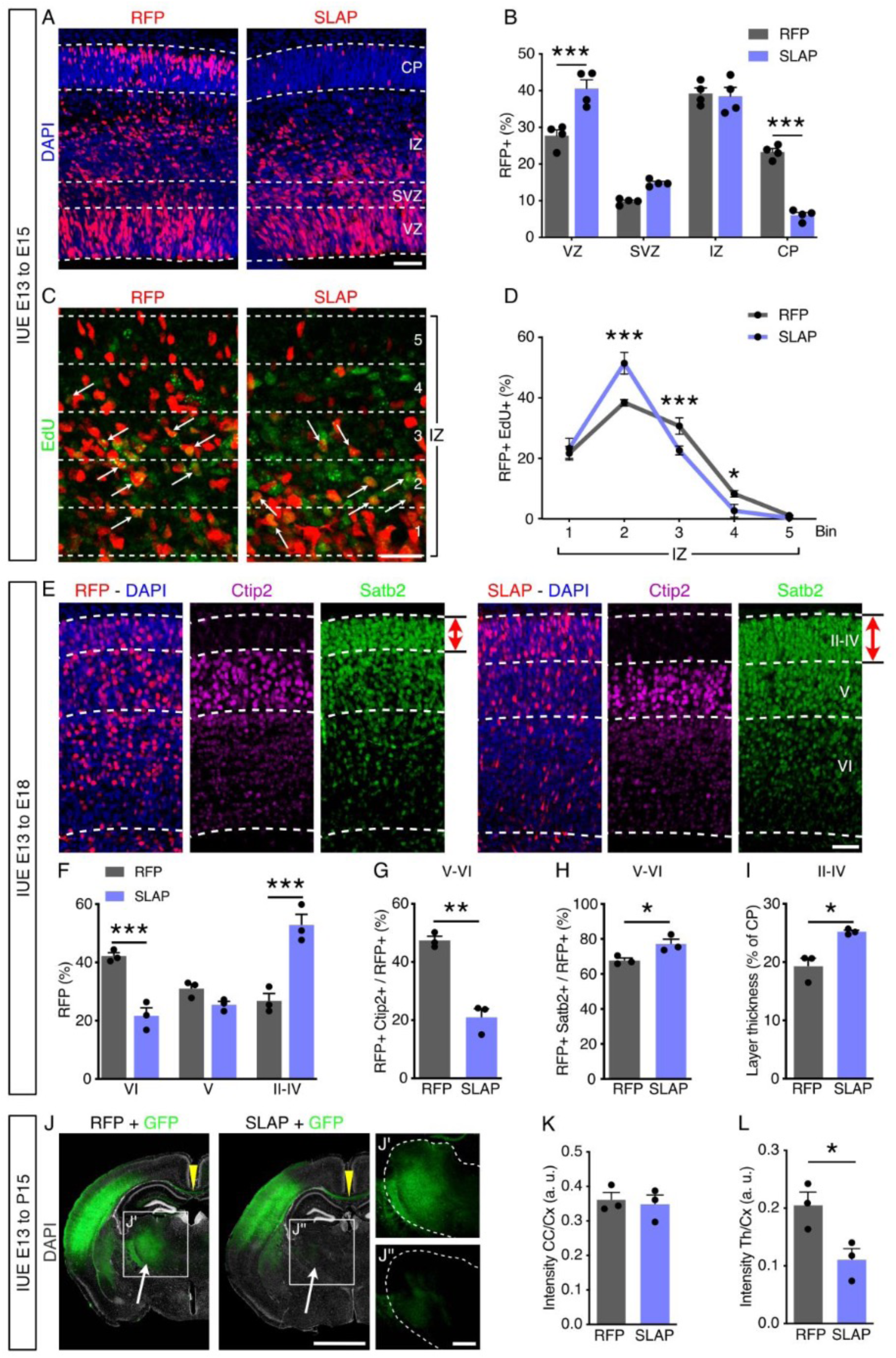
SLAP overexpression alters cortical layering. Fluorescence pictures (A, C, E and J) and quantifications (B, D, F-I, K and L) of coronal sections of the mouse brain from embryos electroporated at E13, and immunolabeled for EdU, Ctip2 or Satb2 (as indicated). Note the changed proportion of EdU+ at E15 (EdU pulse 24 h before sacrifice) across bins of the IZ (C and D), distribution and abundance of RFP+ neurons across cortical layers (II-VI, as indicated) at E18 (E-I), and GFP+ projection to the corpus callosum (CC, arrowheads) or thalamus (Th, arrows) relative to electroporated cortex (Cx) at postnatal day 15 (J-L). Coronal sections are shown in full, including the contralateral non-electroporated hemisphere in Fig. S1E and F. Quantifications are depicted as mean ± SEM stacked lines (D) or bar graphs (B, F-I, K, L). Two-way ANOVA and Bonferroni’s post-hoc test (B, D, F) or a two-tailed Student t test (G-I, K, L) were used to assess significance (* p < 0.05, ** p < 0.01, *** p < 0.001). Scale bars = 50 μm (A, E), 25 μm (C), 2 mm (J), 500 μm (J’’).

First, we evaluated cell cycle exit within the pool of EdU+ electroporated cells that 24 h later became negative for Ki67 finding similar values in control and SLAP targeted cells (RFP = 53.9 ± 3.8 vs. SLAP = 57.0 ± 2.0%, p = 0.5) (Fig. S1A, B). Second, no major difference was observed when assessing apoptosis by cleaved caspase 3 (CC3) immunolabeling (RFP = 0.55 ± 0.04 vs. SLAP = 0.52 ± 0.10 cells/100 μm, p = 0.8) (Fig. S1C, D). Finally, to assess neuronal migration, we analyzed the proportion of RFP+ EdU+ neurons distributed across 5 equal bins across the IZ, and found that control cells localized within the upper bins (3 and 4) to a larger extent than SLAP electroporated cells, which predominantly accumulated in bin 2, closer to the SVZ (RFP = 38.3 ± 0.5 vs. SLAP = 51.1 ± 1.7%, p < 0.001) (Fig. 3C, D).

Furthermore, to assess the effects of SLAP overexpression at later developmental times, we performed electroporation as above and analyzed brains at E18 instead of E15. At this timepoint, we observed that the total proportion of neurons in the CP returned to values overall similar to control (sum of the data shown in Fig. 3F), which is consistent with the transient nature of overexpression by in utero electroporation. However, using the molecular makers Ctip2 and Satb2 to delimit the cortical layers II-IV, V and VI, we found that SLAP overexpressing neurons distributed to a greater extent to the upper (II-IV), and lesser to the deeper (VI), cortical layers compared to control (RFP = 42.1 ± 0.9 vs. SLAP = 21.7 ± 2.2%, p < 0.001) (Fig. 3E, F). In addition to their localization, a decreased proportion of SLAP overexpressing cells were found to be Ctip2+ within deep layer neurons (RFP = 47.3 ± 1.2 vs. SLAP = 20.9 ± 2.4%, p < 0.001) (Fig. 3G) while, conversely, an increased proportion were Satb2+ (Fig. 3H). Notably, this surplus of Satb2+ neurons in SLAP-transfected brains also resulted in thicker cortical layers II-IV (RFP = 19.2 ± 1.1 vs. SLAP = 25.2 ± 0.2% of total CP thickness, p < 0.05) (Fig. 3E red arrows, and I).

Finally, we investigated whether changes in the proportion of upper vs. deeper cortical neurons persisted postnatally and correlated with changes in brain wiring. For these experiments we combined a nuclear-localized RFP with a cytoplasmic GFP during co-electroporation to assess both cell numbers and axonal targeting of projection neurons by the two reporters, respectively. Analyses of postnatal day (P) 15 brains were performed on stereological serial sections showing a consistent targeting of comparable cortical areas, as well as proportion of targeted cells within those areas and targeting almost exclusively restricted to the somatosensory cortex with no other subcortical area being observed to retain fluorescence for the electroporated reporters (Fig. S1E-F).

These analyses at postnatal age confirmed the effects of SLAP overexpression previously observed during development with an increase in the proportion of electroporated cells contributing to upper, at the expense of deeper, cortical layers (Fig. S1G-H). In addition, we quantified mean GFP intensity in the corpus callosum and thalamus and normalized each for the value measured in the targeted portion of the cortex as a proxy to assess differences in the wiring of neurons resulting upon SLAP overexpression. While GFP intensity in the corpus callosum was not altered in SLAP electroporated brains, its intensity in the thalamus was significantly reduced (RFP = 0.20 ± 0.01 vs. SLAP = 0.11 ± 0.01 AU, p < 0.05) (Fig. 3J-L). Given that cortico-thalamic projections derive selectively from deep-layer neurons, these results are consistent with the observed reduction in deep-layer neurons observed at E18 (Fig. 3E-I) (Lodato and Arlotta, 2015), In sum, these results indicated that SLAP overexpression had consequences beyond nuclear integrity and neuronal migration, including effects on neuronal specification, cortical layering and connectivity.

### Knock-down of SLAP expression recapitulates changes in nuclear morphology and cell distribution across cortical layers

Given that artificial overexpression of any gene may result in effects that do not necessarily reflect the physiological function of the endogenous gene, we next attempted to corroborate the phenotypes observed upon SLAP overexpression by the converse manipulation of Cas9-mediated knock-down. To this aim, we identified a 20 bp target sequence within exon 2 of SLAP for CRISPR deletion upon co-electroporation with Ribonucleoprotein complex formed with Cas9 protein and an appropriate gRNA (Fig. 4A) together with RFP reporter plasmid.

**Figure 4.**
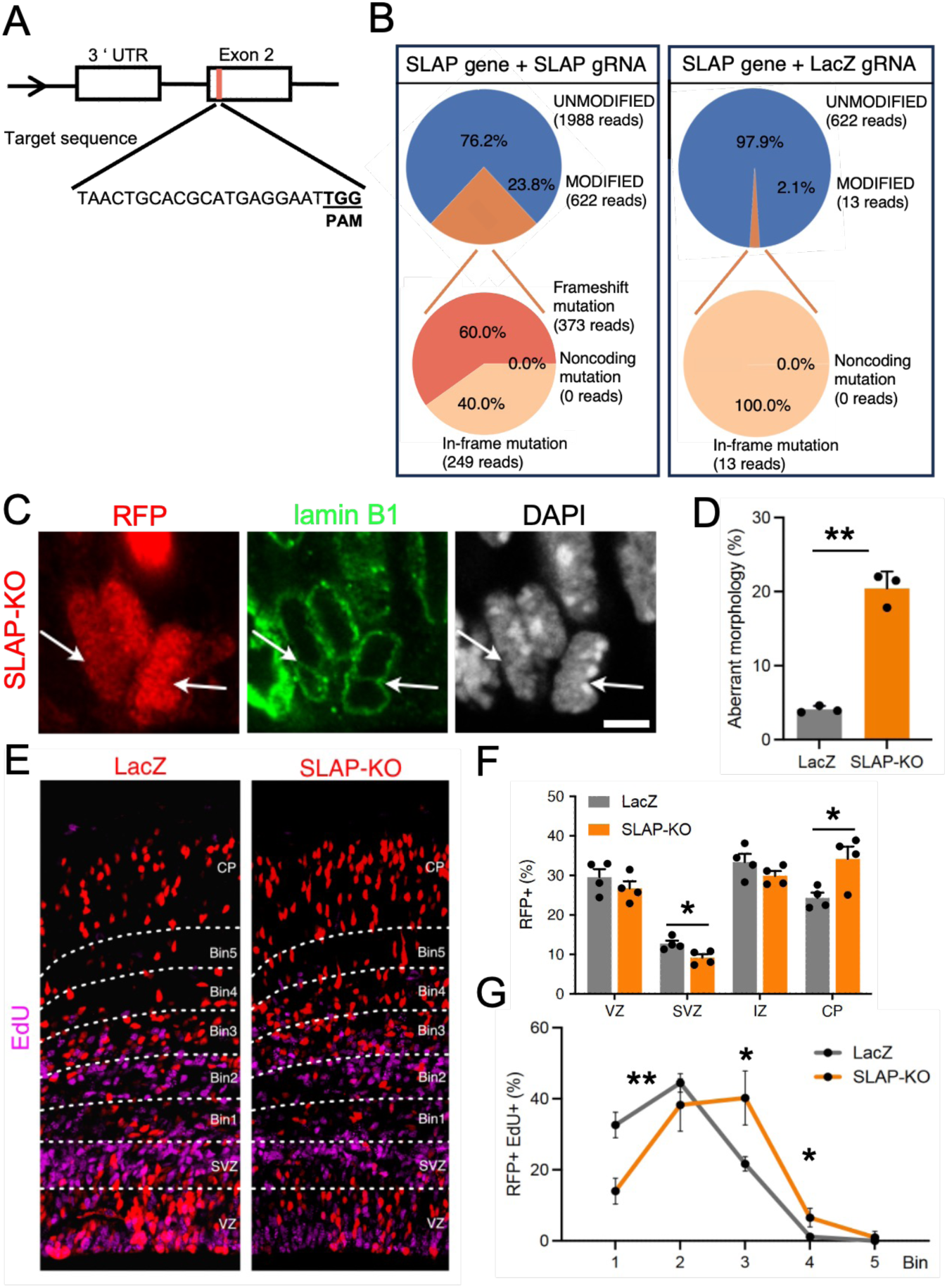
SLAP Knock-down recapitulates overexpression phenotypes. (A) Illustration of the mouse SLAP locus and targeted exon 2 (orange) including gRNA sequence. The protospacer adjacent motif (PAM) is highlighted in bold. (B) Gene modifications observed upon CRISPR/Cas9-mediated mutagenesis using SLAP gRNA or LacZ as control. Panels show the percentage of mutated sequences (upper; orange), frame-shift mutations (lower, dark orange) or unmodified (blue). (C and D) Images and quantification of electroporated apical progenitors counterstained with lamin B1 scored for nuclear abnormalities (arrows). (E-G). Fluorescence pictures (E) and quantifications (F, G) of coronal sections of the mouse brain from embryos electroporated at E13 immunolabeled for EdU (magenta) two days later. Note the change in proportion of EdU+/RFP+ cells (EdU pulse 24 h before sacrifice) across bins of the IZ (G). Quantifications are depicted as mean ± SEM bar graphs (D, F) or stacked lines (G). Two-tailed Student t test (D) or a two-way ANOVA and Bonferroni’s post-hoc test (F, G) were used to assess significance (* p < 0.05, ** p < 0.01). Scale bars = 5 mm (C), 50 mm (E).

Assessment of targeting efficiency and specificity was performed upon in utero electroporation at E13 followed by FAC-sorting of RFP+ cells 48 h later and quantitative sequencing of the SLAP targeted locus and the 5 most similar potential off-target sequences found in the mouse genome (Fig. 4B). Showing the specificity of our approach, none of the two most similar, off-target sequences in the mouse genome showed any significant frame-shift mutation in the SLAP locus (< 0.05%; data not shown) while, conversely, SLAP locus showed a frequency of targeting by 23.8% with frame-mutations occurring in a total of 14.3% of the loci sequenced Fig. 4B). We next investigated whether such levels of SLAP knock-down, despite a somehow relatively low efficacy, was nevertheless sufficient to recapitulate at least some of the results above upon SLAP overexpression on of i) nuclear morphology, ii) proportion of PH3+ mitotic apical progenitors, iii) cell distribution across cortical layers and iv) neuronal migration starting at E15, as described above, upon Cas9 electroporation with SLAP gRNA or LacZ gRNA as negative control.

Intriguingly, some of the effects observed upon knock-down of SLAP expression were similar to those observed upon its overexpression and resulting in both i) aberrant nuclear morphology and ii) increase in PH3+ cells lining the apical border (Fig. 4C and D). In contrast, and as intuitively expected by a converse manipulation, both iii) distribution of targeted cells and iv) neuronal migration showed opposite effects to those observed upon overexpression with an increase in RFP+ cells in the CP and accumulation in upper bins 3-4 upon BrdU neuronal birth-dating at E14 (Fig. 4E-G). Other effects of SLAP knock-down were not assessed or could not be observed (e.g. increase in asymmetric cell divisions or change in the proportion of Tbr2+ cells in the VZ, respectively).

In essence, many of the effects triggered by SLAP knock-down either reproduced or, conversely, mirrored those observed upon its overexpression, despite the fact that Cas9 frame-shift mutations were only observed in 14.3% of loci and, in addition, SLAP protein persisted upon Cas9 targeting. Nevertheless, these results on nuclear morphology, fraction of mitotic cells and neuronal behaviour strengthened our confidence on the biological relevance of our SLAP manipulations, either upon overexpression or knock-down. Hence, we went on to assess the longer-term effects of SLAP overexpression in postnatal mice with specific focus on potential behavioral effects.

### SLAP overexpression during development alters exploratory behaviour in young mice

Given that cognitive deficits resulting from neurodevelopmental disordes are often detectable at an early age (Parenti et al., 2020; Thapar et al., 2017), we next sought to evaluate whether SLAP overexpression induced any observable effect in the behavior of early postnatal mice. To this aim, mouse embryos were electroporated at E13 as described above and analyzed at P15. At this age, juvenile animals begin to leave their nest to explore their environment (Brust et al., 2015). To quantify this behavior, we performed a bedding test in which a box identical to their housing cage was divided into three areas: one with soiled bedding from their home cage, another with clean bedding and an empty area in between (Fig. 5A). One animal at the time was placed in the middle area and recorded for 5 min to analyze locomotion, digging, grooming and rearing.

**Figure 5.**
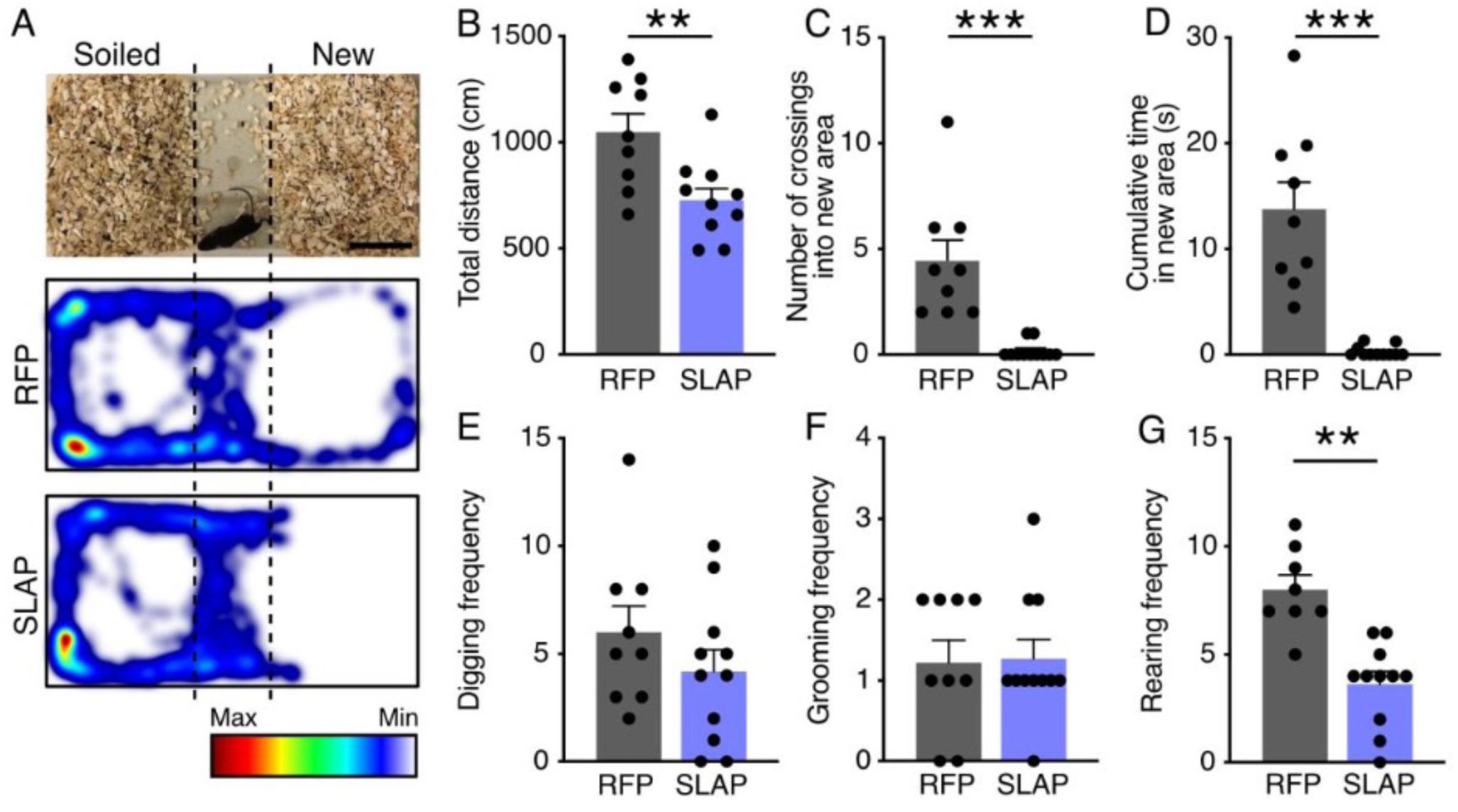
SLAP overexpression reduces exploratory behavior in young animals. (A) Image of the bedding test and representative heatmaps depicting cumulative localization of mice electroporated with RFP or SLAP. Total distance, number of crossings, cumulative time spent in the new area, digging, grooming, and rearing frequency are represented in panels B to G, respectively. Quantifications are depicted as mean ± SEM bar graphs. Two-tailed Student t tests were used to assess significance (** p < 0.01, *** p < 0.001). Scale bar = 5 cm (A).

We found that both RFP and SLAP electroporated animals had no apparent locomotor impediment and identified their home cage bedding reflected by their natural preference to spend most of the time towards that side of the box (Fig. 5A). Nevertheless, SLAP-electroporated animals overall covered less distance and exhibited a greater, and almost complete avoidance, of the new area relative to control mice as assessed by both number of crossings, and time spent, into the home cage bedding area (Fig. 5B-D). While digging and grooming were comparable in both groups of animals, rearing frequency decreased (RFP = 8.0 ± 0.6 vs. SLAP = 3.6 ± 0.5 events in 5 min, p < 0.01) (Fig. 5E-G), again supporting a reduced exploratory behavior in SLAP manipulated animals.

Altogether, these data show the importance of regulating SLAP expression levels during brain development and are reminiscent of manipulations that increased the proportion of upper versus deeper cortical layer neurons related to neurodevelopmental and psychiatric disorders (Falcone et al., 2021; Fang et al., 2014).

## DISCUSSION

As an emerging concept in the field, the NE appears to have more dynamic and complex functions than merely isolating the genome from the cytoplasm. This may apply to all tissues in general (Kalukula et al., 2022) and the central nervous system in particular (Mestres et al., 2022). In addition to control chromatin architecture and provide a mechanical scaffold for nuclear translocation, our study shows that the NE plays further roles in neural stem cell fate, neuronal specification and cortical layer formation, whose impairments can lead to neurodevelopmental disorders.

SLAP overexpression triggered nuclear abnormalities such as invaginations, micronuclei, and ruptures that are hallmarks of several human pathologies ranging from muscle diseases, to cancer and senescence (Kalukula et al., 2022). At the cellular level, nuclear deformations often correlate with defects in nucleokinesis, nucleocytoplasmic traffic, genome instability and DNA damage (Kalukula et al., 2022). It is worth mentioning that in our study, SLAP overexpression triggered changes in nuclear structure and stiffness only in daughter cells, and that nuclear abnormalities were closely associated to a lengthening of metaphase. In line with this, our observations on nuclear morphology defects and proportion of mitotic cells primarily concerned apical progenitors in the VZ more than basal progenitors in the SVZ. While similar results were also observed upon SLAP knock-down, our study could not address the causal relationship between nuclear abnormalities, asymnmetric cell division and cortical layering defects. Remains the fact that metaphase lengthening, changes in tissue stiffness, asymmetric cell division, neurogenic commitment and layering are known to be causally linked (Konno et al., 2008; Noctor et al., 2004; Pilaz et al., 2016).

Until now, SLAP was only associated with lamins and its function awaited clarification (Roux et al., 2012). Although not investigated in our study, we noted that bioinformatic assessment of protein homology (Söding et al., 2005) highlighted a GCN5-related acetyltransferase domain embedded within in the N-terminal half of SLAP (residues 120-230). In turn, this raises the possibility that SLAP is implicated in the acetylation of either lamins, histones or both. In this context, while lamins hyperacetylation did not trigger any major phenotype on nuclear morphology (Karoutas et al., 2019), histone hyperacetylation was described to trigger decreased nuclear stiffness, aberrant nuclear morphology (Stephens et al., 2018), lengthening of pro/metaphase (Cimini et al., 2003) proliferation of neural progenitors and increased cortical neurons (Kerimoglu et al., 2021). The many, close analogies between these previous and our current findings, make it tempting to speculate that the molecular mechanism underlying SLAP function is to modulate histone acetylation influencing brain development and the emergence of neurodevelopmental disorders.

## MATERIALS AND METHODS

### Constructs and Cas9 targetting

SLAP cDNA was generated by RT-PCR from E14 mouse cortex using the following primers, forward: 5’-CAGAATGGCATTCCCTGTGG-3’; reverse: 5’-GGTCAGCTTGGCTTTCTTCT-3’. Amplified fragments were inserted into a pCMS-GFP or pDSV-RFP backbone as previously described (Aprea et al., 2013; Artegiani et al., 2015), and validated by sequencing (Eurofins). Ectopic SLAP was under a cytomegalovirus (pCMS-GFP) or a simian virus 40 (pDSV-RFP) promoter vector with fluorescent reporters (GFP or RFP) under independent SV40 promoters. The sequence of mouse SLAP exon 2 was analyzed for CRISPR/Cas9 target sites by CRISPOR (http://crispor.tefor.net) and gRNA with the highest score and one with the least off-target selected. crRNAfor target sequence, universal tracrRNA (#1072533), and 500 μg HiFi Cas9 nuclease were obtained from IDT (#1081061695) and complex formed in 1:1 (100 μM) molar ratio in Nuclease-free duplex buffer (IDT #11-05-01-03) by incubating at 95°C for 5 minutes. RNP complex formed freshly each time before in-utero electroporation. Embryos were harvested 48 h post-electroporation and dissected brains were treated with Neural Tissue Dissociation Kit P (Macs Miltenyi Biotec, 130-092-628). Genomic DNA from FAC-sorted RFP+ cells upon in-utero electroporation was obtained by Qiagen QIAmp DNA Minikit (# 157050681) followed by PCR amplification of the target regions for 2^nd^ PCR next-generation sequencing (Eurofins). Adaptor sequences were added at the 5’ends of the target PCR primers (Forward Primer: ACACTCTTTCCCTACACGACGCTCTTCCGATCT, Reverse Primer:

GACTGGAGTTCAGACGTGTGCTCTTCCGATCT). NGS data analysis was performed using CRISPResso including two-paired end data merging and indel statistics to assess gRNA modifications on the target site.

### Cell culture

HEK293 cells were seeded onto glass coverslips (Marienfeld, cat. # 0111520) or glass-bottom dishes (Greiner bio-one, cat. # 627870) coated with poly-D lysin and laminin, and kept in DMEM (Thermo Fisher, cat. # 31966-047) supplemented with 10% fetal bovine serum (Gibco, cat. # 10270106) and 1% penicillin/streptomycin. Cells were used between passages 5 and 15, and regularly checked for mycoplasma contamination. For transfection, a 3:1 mix was prepared with polyethylenimine (Sigma, cat. # 408727), DNA added and cells were either fixed or imaged 48 h later.

### In utero electroporation

Wildtype C57BL/6J (Janvier) mice were used with the morning of the vaginal plug defined to as E0. At E13 mice were deeply anesthetized with isoflurane, their uterine horns exposed and approximately 2 μl of plasmid (2 μg/μl with 0.01% of Fast Green) injected into the lateral ventricles. Cas9/gRNA Ribonucleoprotein complex constructs were prepared and in utero electroporated as previously described (Kalebic, et al., 2016). Electrodes were placed adjacent to the embryo head with the anode facing the injection site and 6 pulses of 30 V for 5 ms each delivered using an electroporator (BTX ECM830). The uterus was immediately relocated within the abdominal cavity and the muscular walls and overlying skin sutured independently. Eventually, mice were administered with a single intraperitoneal dose of EdU (4 mg/kg) 24 h after IUE and 24 h before sacrifice. Mice were sacrificed at the indicated time-points by cervical dislocation. Embryo brains were dissected and fixed overnight by immersion in 4% paraformaldehyde (PFA) while postnatal brains were perfused with PBS, 4% PFA, and postfixed in 4% PFA for 2 h. Animal procedures were approved by local authorities (TVV 16/2018).

### Immunochemistry

For immunocytochemistry, cells were fixed with pre-warmed (37 °C) 4% PFA for 15 min; except when using lamin B1 antibodies that required fixation in ice cold methanol with 5% acetic acid for 5 min at – 20 °C. Permeabilization was performed with PBS 1% triton X-100 for 5 min, and blocking with in 5% BSA for 1 h. Either primary (Supp. Table 1) or Alexa-dye conjugated secondary antibodies were incubated in 3% BSA for 1 h at RT. Free-floating coronal brain slices (40 μm thick) were permeabilized and blocked for 1 h at RT in blocking buffer (PBS 0.1% Triton X-100 and 5% donkey serum), and primary (Supp. Table 1) and Alexa-dye conjugated secondary antibodies (1:500, Molecular Probes) incubated at 4 °C in blocking buffer. DAPI was used to counterstain nuclei and the Click-it reaction kit used to reveal EdU (Invitrogen, cat. # C10340).

### Light microscopy and quantifications

Images were acquired using a confocal (LSM 780) microscope or an Axioscan 7 (Carl Zeiss), tiles stitched in Zen and maximal intensity projections quantified using ImageJ (NIH) or Affinity Photo (Serif). Nuclei were scored as aberrant when exhibited micronuclei (condensed fragments of chromatin disconnected from the nucleus), invaginations, blebs, or nuclear envelope ruptures with chromatin scattered into the cytoplasm. Mitotic cleavage orientation and fluorescence intensity were measured with ImageJ. For time-lapse imaging, transfected cells were incubated with SiR-DNA (Tebu-Bio, cat. # SC007) for 2 h prior to imaging to label DNA and a spinning disc microscope (Andor DragonFly) used at 37 °C and 5% CO_2_ with a time resolution of 3 min over a period of ca. 9 h.

For each field, two channels (488 and 633 nm) and about 10 planes were acquired every 3 μm compiled into a maximal intensity projection for analysis. Mitotic phases were defined based of chromatin morphology, dynamics and condensation as follows: prophase, initial chromosome condensation; prometaphase, nuclear envelope disassembly and maximal chromatin condensation; metaphase, chromatin alignment into the metaphase plate; anaphase, sister chromatid separation; telophase, chromatin decondensation.

### Electron microscopy and 3D reconstruction

Electroporated brain sections were immunogold labeled using Nanogold and silver enhancement (Kurth et al., 2010). In brief, the samples were incubated in blocking buffer (PBS 20% goat serum and 0.1% saponin) for 2 h at RT, followed by anti-GPF primary antibody incubation in blocking buffer for 2 days at RT, and anti-rabbit Nanogold (Fab-fragments, Nanoprobes, 1:50) overnight at RT. Samples were postfixed in 1% glutaraldehyde, silver enhanced using the SE-Kit (Aurion), followed by postfixation in 1% osmium tetroxide (1 h on ice), *en bloc* contrasting with 1% uranyl acetate (1 h, on ice), and dehydration in a graded series of ethanol (from 30 to 100%). The samples were infiltrated in Epon substitute EMbed 812, flat embedded on the surface of an empty epon dummy block, and cured overnight at 65 °C. Series of 70 nm sections were prepared using a diamond knife (Diatome) and a Leica UC6 ultramicrotome. The sections were mounted on formvar coated slot grids, and contrasted with 4% uranyl acetate. Serial sections were imaged with a Jeol JEM1400Plus transmission electron microscope running at 80kV acceleration voltage and equipped with a Ruby digital camera (Jeol). For the 3D-reconstruction, series of images of the same cell were manually pre-aligned in Affinity followed by a linear stack alignment in ImageJ, and the nucleus segmented. Signal intensity was processed to binary images, and a 3D gaussian blur was applied (0.5 pixels for all 3 orthogonal axes). A 3D surface render was created with the 3D Viewer tool in ImageJ.

### Atomic force microscopy

Control and SLAP overexpressing HEK cells were seeded 24 h before mechanical probing by AFM onto glass bottom dishes (WPI, Fluorodish). Thirty min before starting the AFM indentation tests, cell culture medium was exchanged to CO_2_-independent medium supplemented with 0.5 µM Cytochalasin D for F-actin depolymerization, and cells were kept at 37 °C using a petridish heater. Indentation tests were conducted using a Nanowizard 4 (JPK Instruments/Bruker) mounted on a Zeiss Axio Observer D1 (Zeiss) microscope. Qpbio cantilevers (nominal Spring constant 0.03-0.09 N/m, Nanosensors) were calibrated before experiments using the thermal noise method using built-in procedures of the scanning probe software. Transfected cells were identified by epifluorescence without considering nuclear morphology as a parameter for the evaluation of stiffness. Then, the cantilever was positioned above the nucleus of a single cell and lowerd at a speed of 5 µm/s until a force of 3 nN was reached. Typically, maximal indentation depths of > 4 µm were thereby reached. In each experiment 60-90 cells were probed. Force-distance curves were processed using the AFM data processing software (JPK DP, JPK Instruments/Bruker), fitting indentation part of the approach curve within a range of 2-4 µm to the Sneddon/Hertz model (half-angle of cone 22.5°, conical indenter). A poisson ratio of 0.5 was assumed.

### Behavior test

Mice exploration was assessed with a bedding test at P14, after habituation to the testing room (15 min) and 3 h before the start of the dark cycle. The test box was identical to their housing cage, subdivided into three areas of which two identical contained 3 day old home cage, or clean, bedding separated by 5 cm space with no bedding in between. The test box and clean bedding were replaced for each individual animal tested. At the beginning of the test, mice were located in the empty center and then recorded for five minutes. Walked distance, crossings areas and time spent in each area were calculated by tracking the body’s center using Ethovision (Noldus). In addition, digging, grooming, and rearing were scored by an experimenter blind to the manipulation.

### Statistical analyses

Quantification of cell types and morphometric analyses were performed on at least three independent biological replicates, and depicted as means ± SEM. Statistical tests were performed using Prism9 (GraphPad). Details on statistical tests and significance are indicated in each figure legend.

## Acknowledgements

We are thankful to Drs. Tomohisa Toda and Mareike Albert for helpful discussion. We are also grateful with the light and electron microscopy (CRTD) and animal (CRTD and MPI-CBG) facilities for their professional assistance, Simone Massalini and Melanie Jäger for excellent technical support, and members of the lab for feedback throughout the project.

## Author contributions

IM and FC conceived the project, analyzed the data and wrote the manuscript. AA performed Cas9 experiments supported by LABC. IM performed all experiments with the assistance of SA, KS, ME, MP. NB conceived behavioral experiments. JCE and AT performed atomic force microscopy, MY supported analyses.

---

**Supplementary Figure 1 (related to Fig. 3).**
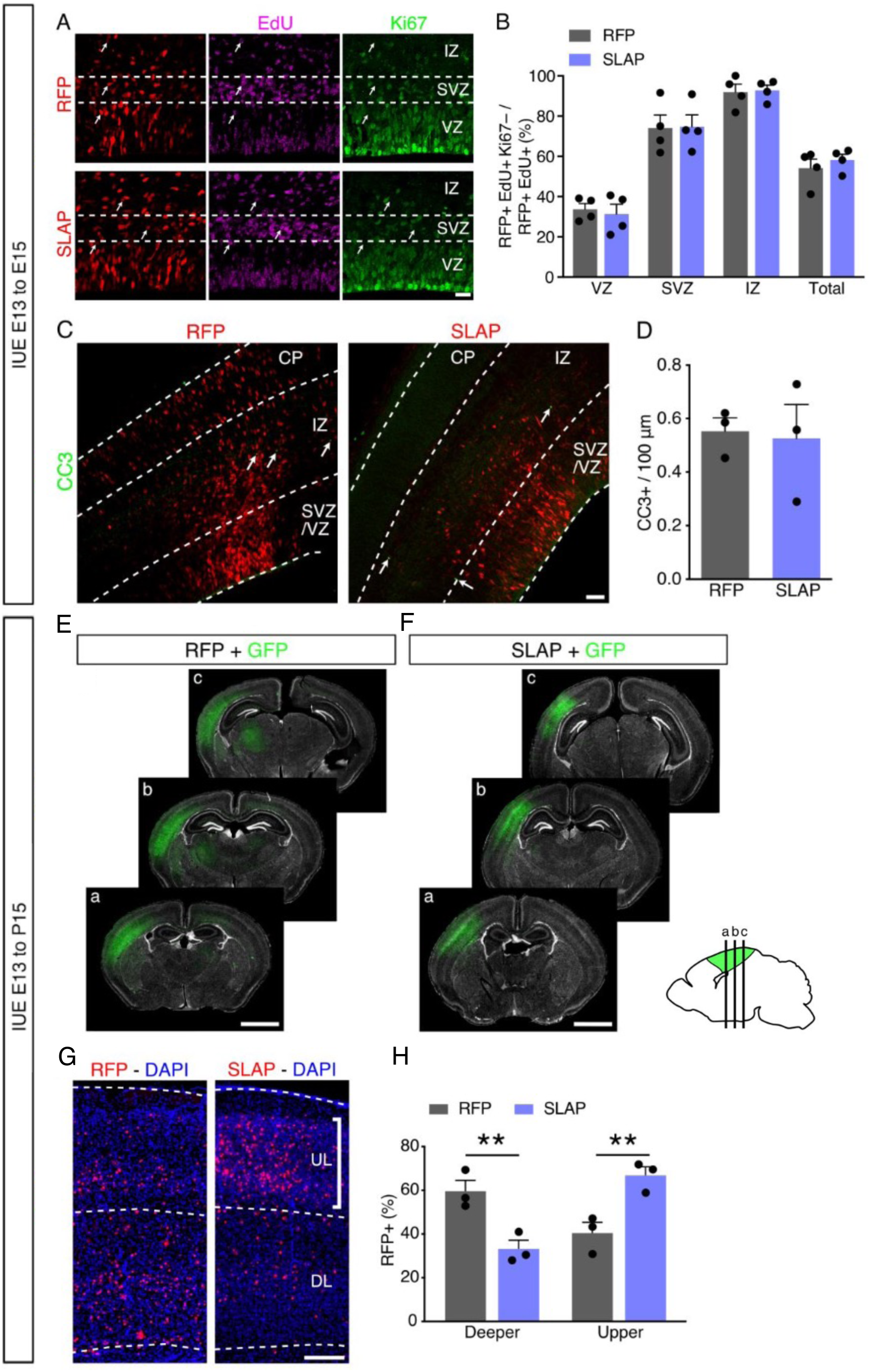
Layering defectes of SLAP overexpression without alterations in cell cycle exit or apoptosis. (A-D) Fluorescence pictures (A and C) of E15 mouse brains electroporated at E13 followed by immunolabeling for markers as indicated (EdU administered 24 h before sacrifice) used to evaluate cell cycle exit (B) or apoptosis (D). (E-F) Representative serial sections of P15 brains electroporated at E13 with GFP (green) together with RFP or SLAP vectors (not whown) and showing targeting of the somatosensory cortex but not subcortical areas. (G and H) Fuorescence pictures of P15 mice electroporated at E13 (G) and quantifications of proportion of of RFP electroporated cells within the upper (UL) and deeper (DL) cortical layers (H). Quantifications are depicted as mean ± SEM bar graphs, Two-tailed Student t test was used to assess significance (** p < 0.01). Scale bars = 25 μm (A), 50 μm (C), 200 μm (G), 2 mm (E, F)

**Supplementary Movie 1.** Related to Figure 1. Time-lapse imaging of HEK293 cells 48 h after transfection with SLAP (green). Gray channel shows chromatin. White arrows point to the transfected cell (right) or an untransfected cell (left) going through mitosis. Nuclei within one daughter (red arrows) and a micronucleus (arrowhead) are indicated. Time is shown as h:min. Scale bar 20 μm.

**Supplementary Movie 2.** Related to Figure 2. Nuclear 3D reconstruction of a SLAP electroporated apical progenitor in the mouse E15 brain.

**Supplementary Table 1.**
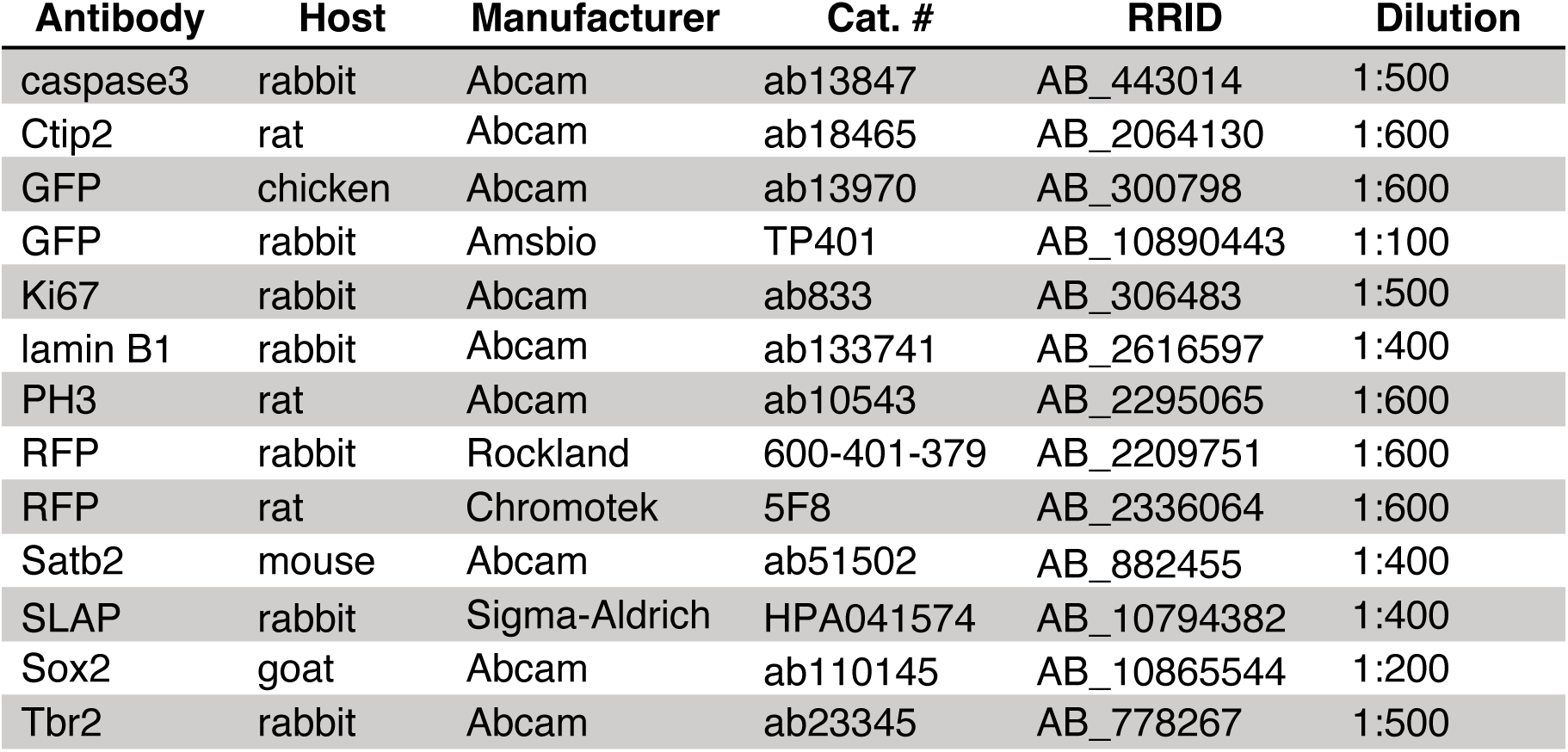
List of antibodies used in this study.

